# Psychological scales in the brain: Trait-linked questionnaire items evoke similar neural patterns in the mPFC

**DOI:** 10.1101/2025.08.21.671650

**Authors:** Keise Izuma, Ayahito Ito, Kazuki Yoshida, Ryuta Aoki

**Affiliations:** School of Economics & Management, Kochi University of Technology, Kochi 780-8515, Japan; Research Center for Mind, Brain, and Behavior, Kochi University of Technology, Kochi 780-8515, Japan; School of Psychology, University of Southampton, Southampton, SO17 1BJ, UK; Graduate School of Education, Tohoku University, Sendai, 980-8576, Japan; Faculty of Health Sciences, Hokkaido University, Sapporo, 060-0812, Japan; Graduate School of Medicine, Human Brain Research Center, Kyoto University, Kyoto, 606-8507, Japan

**Author notes:** Keise Izuma, School of Economics & Management, Kochi University of Technology, Kochi 780-8515, Japan.

**Keywords:** self-report, self-referential processing, mPFC, default mode, personality assessment, Big Five

## Abstract

Self-report questionnaires are widely used across psychology and related disciplines, yet the cognitive and neural processes underlying how individuals generate responses to such items remain poorly understood. Here, we investigated whether items from the same psychological scale evoke similar neural activation patterns in the medial prefrontal cortex (mPFC), a region consistently implicated in self-referential processing. While undergoing functional magnetic resonance imaging (fMRI), participants completed a self-reference task in which they judged how well 72 personality-related questionnaire items (e.g., from the Big Five, emotion regulation, and well-being scales) described themselves. Using representational similarity analysis (RSA), we found that items from the same scale elicited more similar multivoxel activation patterns in the mPFC compared to items from different scales. This effect was specific to the self-reference task and was not observed during a semantic judgment control task using the same items. Furthermore, the mPFC encoded not only categorical scale membership but also graded psychological similarity among scales, as reflected in inter-scale behavioral correlations. Importantly, these effects remained significant even after controlling for sentence-level semantic similarity using multiple regression RSA, indicating that the observed neural structure reflects psychological rather than linguistic similarity. These findings suggest that the mPFC integrates internally constructed evidence in a construct-sensitive manner during self-report, and they open new avenues for linking psychological assessment with neural representation. We discuss the implications for understanding self-report as a cognitive process and for future work on neuroimaging-informed scale validation.

**Significance statement:** Self-report questionnaires are widely used across psychology, medicine, and public policy to assess thoughts, feelings, and personality traits. However, little is known about how the brain generates these responses. Using fMRI and a multivariate analysis approach, we found that questionnaire items measuring the same psychological construct evoked similar activity patterns in the medial prefrontal cortex—a region involved in self-reflection. This effect was specific to self-referential judgments and remained significant even after controlling for sentence-level semantic similarity, suggesting that the brain organizes information according to underlying psychological traits rather than linguistic features. These findings offer new insights into how self-report operates at the neural level and may inform future approaches to scale development.

## Introduction

The ability to report one’s own inner thoughts and feelings is a hallmark of the human species. Across the social sciences, self-report has long served as a foundational research tool, offering direct access to individuals’ internal experiences and shaping our understanding of human psychology and behavior. Beyond psychology, self-report remains a primary method of data collection in a wide range of disciplines, including medicine, education, political science, economics, public health, and marketing. Its widespread use reflects its unique capacity to capture subjective experiences that are otherwise inaccessible through external observation alone^1,2^.

Although the psychological processes underlying self-report have been a central topic in psychology^2^, the neural mechanisms supporting this process remain poorly understood. While social and cognitive neuroscience studies have employed a wide range of tasks in which participants report their attitudes, preferences, feelings, and beliefs (often using Likert scales) during brain imaging, these findings have not been synthesized into a broader framework. As a result, our understanding of how the brain generates responses to self-report questions remains limited.

One widely used approach in social neuroscience to investigate self-referential processing is the self-reference task^3,4^, in which individuals are presented with personality trait adjectives and asked to judge whether each adjective describes them—a format similar to personality questionnaires. This task has consistently been shown to activate a network of brain regions, most notably the medial prefrontal cortex (mPFC) and posterior cingulate cortex (PCC)^5,6^. These regions are recognized as core hubs of the brain’s default mode network^7^ and are commonly engaged in tasks that require judgment based on internally constructed representations, such as theory of mind, autobiographical memory, spatial navigation, moral reasoning, and self-related evaluation^8-13^.

Our previous study^14^ demonstrated that, during the self-reference task, individuals— consciously or unconsciously—engaged in multiple cognitive processes to gather the information necessary to evaluate whether a personality trait described them. This multifaceted evaluative process was reflected in distinctive activation patterns within the mPFC, suggesting that the mPFC plays a central role in integrating internally constructed representations to support self-referential judgments.

Building on this finding, the present study aims to further investigate the integrative function of the mPFC by examining whether its activation patterns during self-report questionnaire responding reflect the psychological constructs measured by each item. While undergoing functional magnetic resonance imaging (fMRI), participants completed several psychological trait scales. As a control task, participants also rated the social desirability of the same questionnaire items, allowing us to distinguish neural activity associated with self-referential evaluation from that involved in more general semantic or normative judgments. Whereas the self-reference task required participants to retrieve and integrate personally relevant information to evaluate whether each item described them, the desirability judgment task emphasized reasoning based on socially shared norms and expectations (i.e., externally guided decision-making; see ^8^).

As the present study is, to our knowledge, the first neuroimaging investigation of questionnaire responding using a multivariate analysis approach, we selected the Big Five personality dimensions as our primary experimental stimuli due to their central role in personality psychology and broad applicability across research domains. The Big Five framework is among the most widely used and empirically supported models of personality, offering a parsimonious yet comprehensive taxonomy of trait-like individual differences^15,16^. Its enduring influence is evident not only in its widespread use in psychological assessment, but also in its integration into related fields such as behavioral genetics^17,18^, developmental psychology^19,20^, and organizational behavior^21,22^. Recent work further supports the Big Five’s utility as an organizing framework for psychological trait constructs: Bainbridge et al.^23^, for example, demonstrated that many of commonly used stand-alone trait measures—including constructs such as self-esteem, grit, and empathy—can be meaningfully located within the Big Five space, often correlating as strongly with Big Five domains as with their originally associated scales. This supports the use of Big Five items for probing the neural representation of psychological traits, as they provide both theoretical clarity and empirical breadth. Accordingly, by focusing on the Big Five, the present study leverages a widely accepted framework to examine how psychological constructs are instantiated in brain activation patterns during self-referential evaluation.

To analyze the neural data, we applied representational similarity analysis (RSA)^24^, a multivariate approach that tests whether local patterns of neural activity reflect specific psychological structures. In the present study, we asked whether multivoxel patterns in the mPFC reflect item-level similarity within or across Big Five trait domains. By comparing these neural patterns to model representational similarity structures—defined theoretically or derived from behavioral data—we sought to clarify the type of information encoded in the mPFC during questionnaire responding.

Tourangeau et al.^2^ proposed an influential model of the cognitive processes involved in self-report, identifying four key stages: comprehension, retrieval, judgment, and response. First, respondents must understand the question (comprehension stage). Second, they must recall relevant information from memory (retrieval stage), such as episodic memories (e.g., “When was the last time I felt anxious?”) or more abstract self-knowledge (e.g., “Am I generally an anxious person?”). Third, once information is retrieved, respondents must evaluate it and formulate a response (judgment stage), such as summarizing across multiple memories or integrating conflicting information. Finally, they must map this internal judgment onto the response options provided (e.g., a Likert scale) (response stage). Whereas the first and last stages are shared across the self-reference and control (desirability judgment) tasks, the retrieval and judgment stages in the self-reference task uniquely involve constructing and evaluating internally generated representations. Accordingly, we hypothesized that to the extent that well-constructed psychological scales prompt individuals to access similar kinds of internal information, items assessing the same trait dimension would evoke similar multivoxel activation patterns in the mPFC, reflecting a shared evidence integration process.

## Methods

### Participants

We aimed to recruit a minimum of 30 participants based on our previous studies^14,25^ that employed similar self-referential tasks and multivariate analysis approaches. The final sample consisted of 32 participants (14 women, 18 men) aged 18–26 years (M = 22.09, SD = 1.71). A total of 38 right-handed individuals were initially recruited through the university participant pool at Hokkaido University. Data from six participants were excluded due to excessive head motion (>3 mm; one participant), scanner malfunction (four participants; EPI data from these individuals could not be normalized due to low signal quality), and low reliability in self-reference or desirability ratings (one participant). For the reliability exclusion, we calculated within-task correlations across two repetitions of the same ratings. The excluded participant showed a correlation of 0.46 for the self-reference ratings, well below the average (*r* = 0.78) observed in the remaining participants, suggesting low task compliance or instability in self-evaluative responses. This exclusion criterion (i.e., *r* < 0.5) is consistent with that used in our previous study^25^.

All participants reported no history of psychiatric disorders. Written informed consent was obtained from all participants prior to the fMRI session. The study was approved by the ethics committee of Kochi University of Technology as part of a joint research project between Hokkaido University and Kochi University of Technology. Participants received 4,000 Japanese yen as compensation.

### Experimental procedure

#### Experimental stimuli

Experimental stimuli consisted of 72 questionnaire items in total: 60 items from Big Five Inventory-2 (BFI-2)^16^, six items from the Psychological Well-Being (PWB): Purpose in Life scale^26^, and six items from the Emotion Regulation (ER): Reappraisal scale^27^. The five dimensions of the BFI-2 are known to exhibit low intercorrelations, and the two additional scales were included based on prior findings showing that they are not highly correlated with any of the Big Five dimensions^23^. All scales had been previously translated into Japanese and validated with Japanese samples^28-30^. While the original PWB: Purpose in Life scale includes seven items, one item (“Done all there is to do in life”) was excluded, as it showed divergent response patterns from the rest of the scale in a Japanese sample, resulting in a lower Cronbach’s alpha in prior research^28^.

#### fMRI experiment

Prior to the fMRI scan, participants received instructions regarding MRI safety and the tasks they would perform inside the scanner. During the fMRI session, participants were presented with one questionnaire item and a 4-point scale on each trial and completed two tasks: self-reference and semantic judgment (Figure 1; see details below). In the self-reference task, participants were asked to judge how much each item described them. In the semantic judgment task, they rated how desirable each item was. For both tasks, responses were made using a 4-point scale (self-reference: not at all – very much; semantic judgment: very undesirable – very desirable).

**Figure 1.**
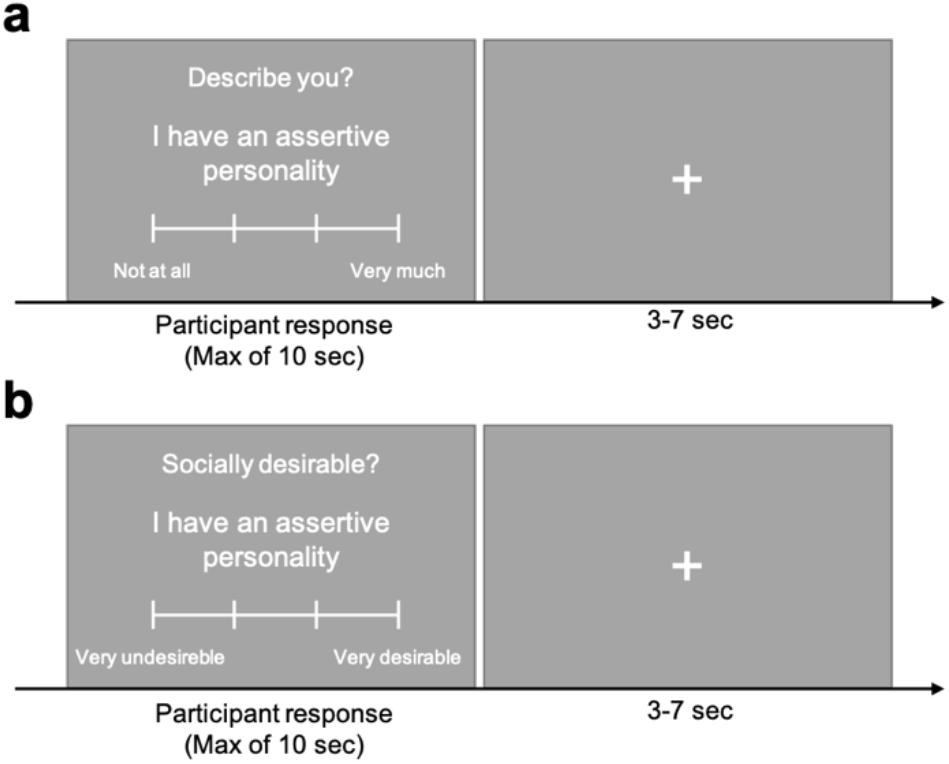
fMRI tasks. (**a**) Self-reference task. (**b**) Semantic judgment task.

Each item remained on the screen until the participant responded, with a maximum response window of 10 seconds. If no response was made within this time limit, a warning message (“Respond faster”) was displayed for 2 seconds. Trials were separated by a jittered inter-trial interval (ITI) ranging from 3 to 7 seconds (mean = 5 seconds). Participants provided their responses using an MRI-compatible button box with four buttons. To prevent systematic associations between specific fingers and response values, the mapping between fingers and rating options was reversed after the fourth run (of eight total runs).

Each of the 72 questionnaire items was presented four times in total—twice for each of the two tasks—so that participants made both self-reference and semantic ratings twice per item. This repetition was implemented to increase the signal-to-noise ratio^31^. However, to prevent participants from simply recalling and repeating their previous responses without thoughtful engagement, we instructed them that inconsistencies across repetitions were acceptable and asked them to respond as if they were encountering each item for the first time on every trial.

Participants completed four runs of each task. Each run consisted of 36 trials, with one questionnaire item presented per trial, and the trial order was randomly determined within each run. The same set of 36 items was presented across both tasks and in each run. Task order was counterbalanced across participants using one of two fixed sequences: ABBAABBA or BAABBAAB. The experimental tasks were programmed using Psychtoolbox (http://psychtoolbox.org/) with MATLAB software (version 2018a, http://www.mathworks.co.uk).

### fMRI data acquisition

We acquired images using a 3-T Siemens MAGNETOM Prisma scanner equipped with a 20-channel head coil. For functional imaging, interleaved T2*-weighted gradient-echo echo planar imaging (EPI) sequences were used to produce 42 contiguous 3.0-mm-thick transversal slices that cover nearly the entire cerebrum (TR = 2,500 ms; TE = 30 ms; flip angle = 90°; field of view = 240 mm; 80 × 80 matrix; voxel size = 3.0 × 3.0 × 3.0 mm). High-resolution anatomical T1-weighted images were also acquired for each participant (TR = 2,300 ms; TE = 2.32 ms; flip angle = 8°; field of view = 240 mm; 256 × 256 matrix; voxel size = 1.0 × 1.0 × 1.0 mm).

### fMRI data preprocessing

We carried out preprocessing and statistical analysis of the fMRI data using SPM12 (Wellcome Department of Imaging Neuroscience), implemented in MATLAB (MathWorks). The first four volumes were discarded before preprocessing and data analyses to allow for T1 equilibration. We conducted preprocessing of the fMRI data with SPM 12’s preproc_fmri.m script starting with realignment of all functional images to a common image. We spatially realigned all images within each run to the first volume of the run using 7th-degree B-spline interpolation and unwarped and corrected for motion artifacts.

The T1-weighted structural image was segmented and normalized into a common stereotactic space (MNI atlas). Subsequently, the normalization parameters were applied to the functional images and resampled them to 3 × 3 × 3 mm^3^ isotropic voxels (i.e., original voxel size was retained) using 7th-degree B-spline interpolation. Following the normalization, the data were spatially smoothed (with a Gaussian kernel of 8 mm FWHM) for the univariate analysis. To preserve fine-grained activation patterns, smoothing was not applied prior to the first-level data analysis for the RSA. Instead, smoothing was applied only at the group level to the RSA-derived maps using a 4 mm FWHM Gaussian kernel, to account for individual variability in brain structure.

### fMRI data analysis

#### Univariate analysis

We used two general linear models (GLMs) to analyze the fMRI data. For the univariate analysis, which aimed to replicate the robust activation of the mPFC during self-referential judgments compared to semantic judgments, we modeled all trials from each condition (self-reference and semantic judgment) with separate regressors. The six head motion parameters were included as nuisance regressors. A contrast image comparing the self-reference and semantic judgment conditions was then created for each participant and submitted to a second-level group analysis (i.e., one-sample t-test across participants).

For the RSA, we estimated a separate first-level GLM in which each questionnaire item was modeled individually (72 items × 2 repetitions × 2 tasks = 288 regressors across 8 runs). Trials with missing responses were modeled as a regressor of no interest, and the six head motion parameters were again included as nuisance regressors. Contrast and t-maps were created for each item in each condition, resulting in a total of 144 maps (72 per task).

Importantly, although each repetition was modeled separately at the first level, for the contrasts used in RSA we combined the two repetitions of each item (assigning a weight of 1 to both repetitions). Thus, each t-map reflected the average activation pattern across repetitions for that item. These t-maps were then used in the subsequent RSA analyses.

Within the mPFC mask (see below), we set statistical threshold at *p* < 0.005 (uncorrected), with a cluster-level threshold of *p* < 0.05 (FWE corrected). We also conducted a whole brain analysis using the same thresholds without the mask; a height threshold at *p* < 0.005 (uncorrected), with a cluster-level threshold of *p* < 0.05 (FWE corrected).

#### Representational similarity analysis (RSA)

##### Neural representational similarity matrix (RSM)

To test whether questionnaire items from the same psychological scale evoke similar activation patterns, we performed RSA using a searchlight approach ^32^. Local patterns of neural activity were extracted from spherical searchlights with a 3-voxel radius, each consisting of up to 123 voxels (fewer near the edges of the brain). For each searchlight, we computed voxel-wise correlations between all possible pairs of the 72 items, resulting in a 72 × 72 neural RSM (Figure 2a). Each cell of the neural RSM represented the correlation in neural activation between a given pair of items within that specific searchlight.

**Figure 2.**
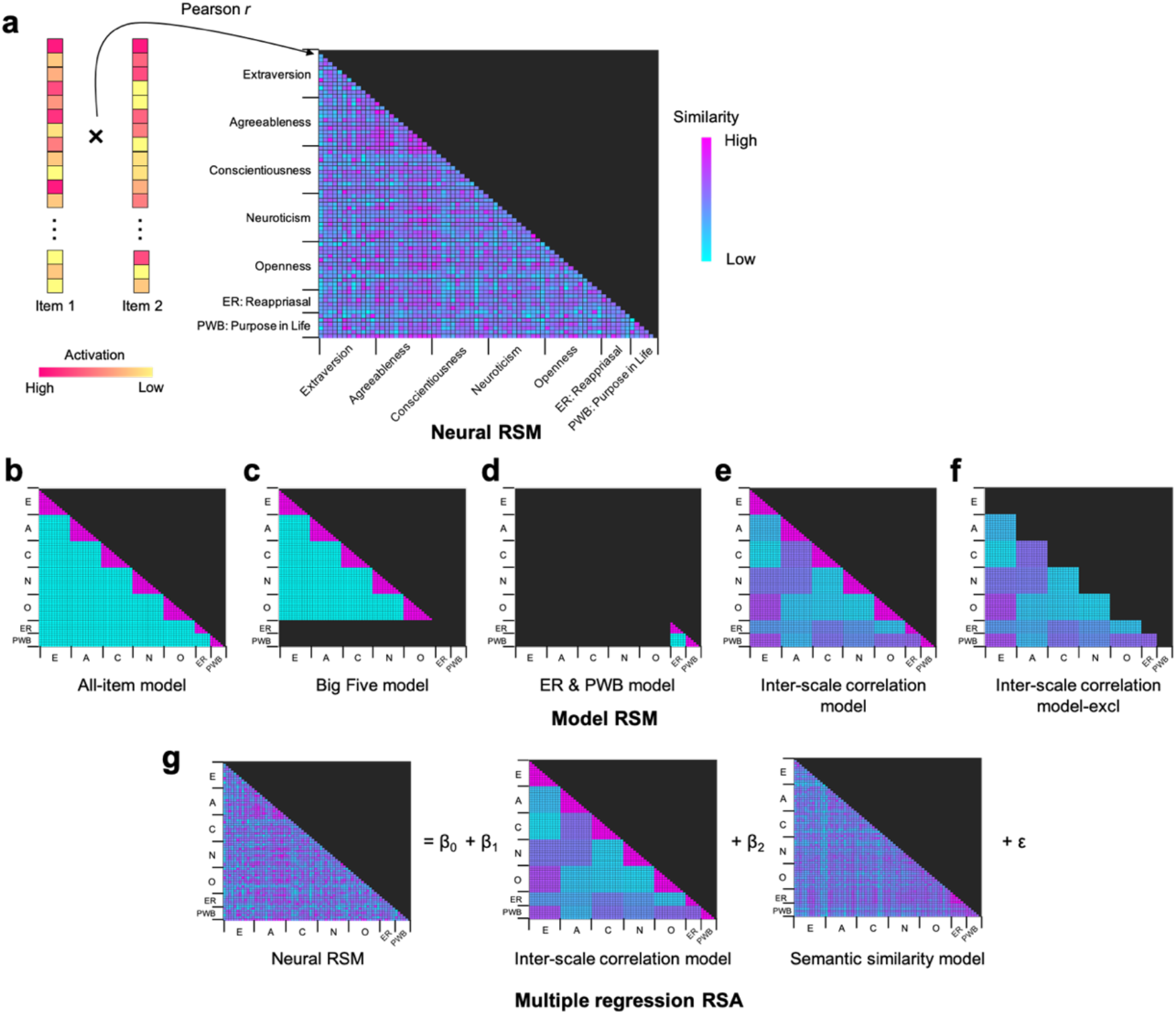
Schematic illustrations of representational similarity analysis (RSA). (**a**) For each participant, a neural representational similarity matrix (RSM) was constructed by computing Pearson correlations between multivoxel activation patterns for all possible pairs of questionnaire items. (**b**) The all-item model included all 72 items from the seven psychological scales. Magenta indicates item pairs coded as similar (1), cyan as dissimilar (0), and black cells were excluded from analysis. (**c**) The Big Five model included only the 60 items from the Big Five Inventory. (**d**) The ER & PWB model included the 12 items from the Emotion Regulation (ER) and Psychological Well-Being (PWB) scales. (**e**) The inter-scale correlation model included all 72 items. Each cell value reflects the absolute value of the correlation coefficient between the two scales to which the items belong (see Figure 3). (**f**) The inter-scale correlation model–excl was identical to the inter-scale correlation model, except that cells corresponding to item pairs from the same scale (i.e., the magenta triangle regions in panel e) were excluded from analysis. (**g**) Neural and model RSMs in the multiple regression RSA. Semantic similarity model shown here is based on Sentence-BERT. E = Extraversion; A = Agreeableness; C = Conscientiousness; N = Neuroticism; O = Openness; ER = Emotion Regulation; PWB = Psychological Well-Being.

##### Model RSM

We tested a total of five model RSMs (Figure 2b-f); (1) All-item model, (2) Big Five model, (3) ER & PWB model, (4) inter-scale correlation model, and (5) inter-scale correlation model-excl. The all-item model (Figure 2b) included all 72 items and tested whether neural responses were more similar for item pairs from the same psychological scale than for those from different scales. The Big Five model (Figure 2c) was identical in structure but included only the 60 items from the Big Five Inventory, isolating effects specific to those dimensions. The ER & PWB model (Figure 2d) included only the 12 items from the Emotion Regulation and Psychological Well-Being scales to examine whether effects observed in the Big Five model were also present in other psychological constructs.

The inter-scale correlation model (Figure 2f) tested whether neural similarity reflected not only category membership (same vs. different scale) but also the graded degree of behavioral correlation between scales—i.e., whether items from scales that are more strongly correlated across individuals elicited more similar activation patterns in the mPFC. To construct this model, we first computed mean ratings for each of the seven scales (five Big Five dimensions, ER, and PWB) for every participant. We then calculated Pearson correlations across participants between each pair of scales, yielding a 7 × 7 inter-scale correlation matrix (reported in Figure 3). This matrix was expanded to the item level by assigning each pair of items the absolute correlation value of their respective scales, resulting in a 72 × 72 model RSM. The inter-scale correlation model–excl was identical to the inter-scale correlation model, except that it excluded all within-scale item pairs (i.e., the cells along the block diagonals of the RSM), allowing us to assess whether between-scale correlations drove the observed neural similarity patterns.

**Figure 3.**
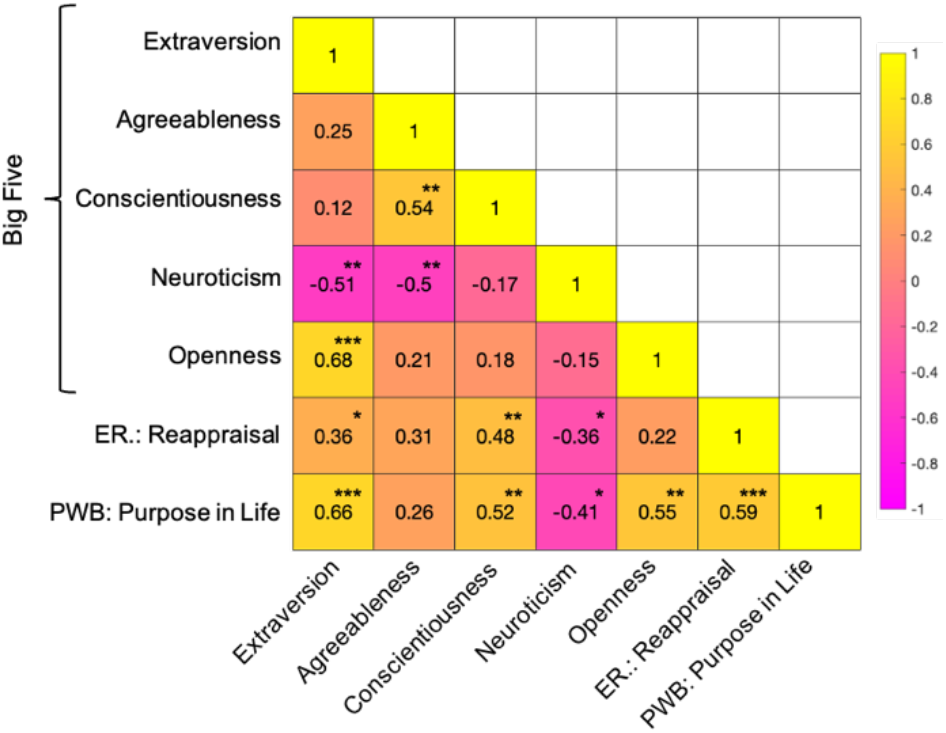
Correlations across the seven scales. *** *p* < 0.001, ** *p* < 0.01, * *p* < 0.05 (two-tailed, uncorrected for multiple comparisons).

The fit between the neural RSM and each model RSM was evaluated by using Kendall’s tau-a for each participant (Nili et al., 2014). We repeated this analysis for every searchlight across the whole brain, resulting in a correlation map for each of the two tasks.

##### Group analysis

As mentioned above, we applied smoothing before the group analysis of the RSA outputs (with a Gaussian kernel of 4-mm FWHM). Following the smoothing, the correlation maps were submitted to the second-level analysis (i.e., one sample t-tests across participants). We applied an mPFC mask to the analysis to restrict the analysis to voxels within the *a priori* mPFC ROI. We created the mask in the same way as in our previous studies^25^ using the WFU PickAtlas toolbox for SPM^33^. The mPFC ROI mask included Frontal_Sup_Medial_L, Frontal_Sup_Medial_R, Frontal_Mid_Orb_L, Frontal_Mid_Orb_R, Rectus_L, Rectus_R, Cingulum_Ant_L, and Cingulum_Ant_R (dilation factor = 2), which we took from the Anatomical Automatic Labeling (AAL) masks implemented in the WFU pickatlas toolbox.

Within the mPFC mask, we set statistical threshold at *p* < 0.005 (uncorrected), with a cluster-level threshold of *p* < 0.05 (FWE corrected). We also conducted the whole brain analysis using the same thresholds without the mPFC mask; a height threshold at *p* < 0.005 (uncorrected), with a cluster-level threshold of *p* < 0.05 (FWE corrected).

#### ROI analysis

##### RSA

To further investigate the role of the mPFC, we ran a ROI analysis. We defined our mPFC ROI in the same way as in our previous studies^14,25^, using Neurosynth (https://neurosynth.org/)^34^. We downloaded an association map (thresholded at *q* < 0.01, False Discovery Rate–corrected) which was generated from a term-based meta-analysis with the term “self-referential” (downloaded October, 2023). The mPFC ROI included 308 voxels.

The RSA was conducted within this ROI. For each participant, we computed the fit between the neural RSM and each model RSM within the mPFC ROI. Statistical significance was assessed using a permutation test in which the order of the 72 questionnaire items was randomly shuffled 1,000 times to generate a null distribution.

##### Multiple regression RSA

To test whether our RSA results held after controlling for semantic similarity among scale items, we conducted a multiple regression RSA in which the mPFC neural RSM derived from activation patterns of the mPFC ROI served as the dependent variable (see Figure 2g). The model included two predictors: the inter-scale correlation model (Figure 2e) and the semantic similarity model, both of which were z-scored prior to analysis. Semantic similarity was first estimated using Sentence-BERT (Reimers & Gurevych, 2019), a transformer-based model that generates sentence embeddings such that semantically similar sentences occupy nearby positions in embedding space. For the Japanese items, we used a pre-trained Japanese BERT model (cl-tohoku/bert-base-japanese-whole-word-masking) implemented within the Sentence-BERT framework. We computed cosine similarity between all possible pairs of questionnaire items and used the resulting 72 × 72 matrix as the semantic similarity model RSM (Figure 2g).

To ensure that our results were not dependent on the embedding method, we repeated the analysis using Google’s Universal Sentence Encoder (USE)^35^, a deep averaging network that encodes sentences into fixed-length vectors optimized for semantic similarity and transfer learning tasks. Specifically, we used the multilingual version of USE, which supports over a dozen languages, including Japanese. The USE-based semantic similarity RSM was substituted into the regression analysis in place of the Sentence-BERT model (Figure 2g). The two semantic similarity RSMs were moderately correlated with one another (*r(2554)* = 0.34, *p* < 0.001), suggesting that they captured overlapping but non-identical semantic structure. Statistical significance was assessed using a permutation test, as described above.

## Results

### Behavioral results

Participants responded within the 10-second time limit on nearly all trials: 99.93% for the self-reference task and 99.87% for the semantic judgment task. The average reaction time (RT) was 2.89 seconds (SD = 0.69) for the self-reference task and 2.85 seconds (SD = 0.74) for the semantic judgment task. This difference was not statistically significant (*t*(31) = 0.93, *p* = 0.36).

Participants rated each of the 72 items on self-descriptiveness and desirability twice each during the fMRI session. Their responses were highly consistent across the two repetitions. The average within-individual correlation for the self-reference task was *r* = 0.79 (SD = 0.11), and for the semantic judgment task, *r* = 0.87 (SD = 0.07). Response consistency was significantly higher in the semantic judgment task than in the self-reference task (paired t-test on Fisher z-transformed values: *t*(31) = 6.01, *p* < 0.001).

We further examined whether participants exhibited a typical self-enhancement tendency ^14,36^—that is, whether more desirable items were more likely to be endorsed as self-descriptive. For each participant, we calculated the Pearson correlation between self-descriptiveness and desirability ratings (using the average of the two ratings per item). A one-sample t-test on Fisher z-transformed correlation coefficients revealed that the average correlation (*r* = 0.38, SD = 0.36) was significantly positive (*t*(31) = 5.90, *p* < 0.001), indicating a reliable self-enhancement effect.

Using the average of the two ratings for each item, we computed Cronbach’s α for each of the seven psychological scales. All scales demonstrated good internal consistency (Cronbach’s α > 0.80), as shown in Table 1. Correlations among the seven scales are presented in Figure 3.

**Table 1.**
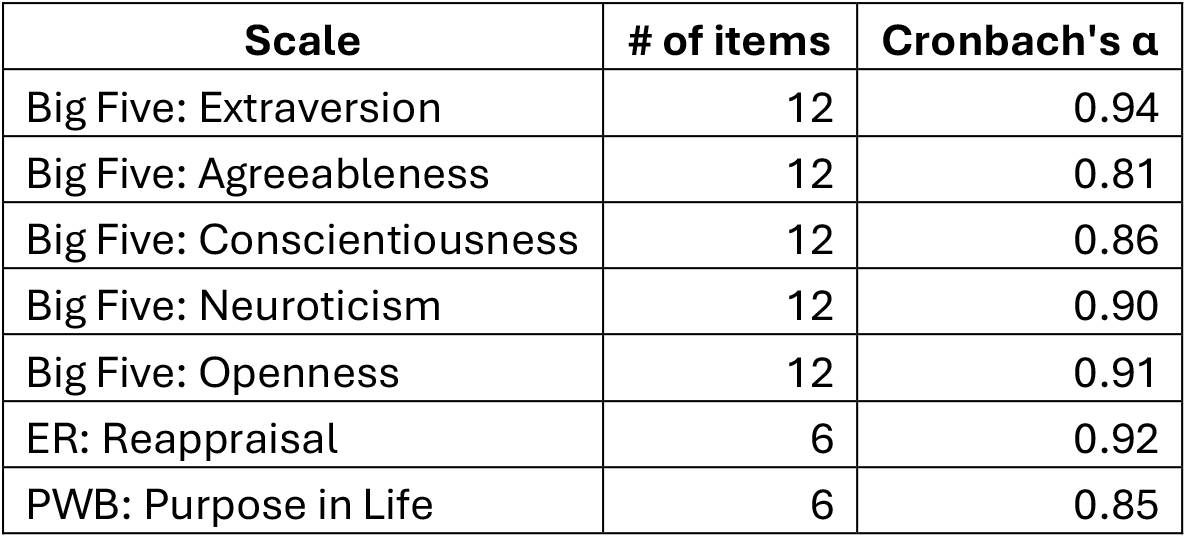
Cronbach’s α for the seven scales.

Finally, we examined the similarity of self-descriptiveness and semantic ratings across all participant pairs for the 72 questionnaire items (see Figure 4). Correlations for the semantic ratings were uniformly positive, ranging from *r* = 0.57 to *r* = 0.93, indicating that participants rated the desirability of items in a similar way—that is, they shared a common understanding of social norms (Figure 4b). In contrast, correlations for self-reference ratings showed greater variability, ranging from *r* = –0.48 to *r* = 0.69, suggesting that participants differed in how they judged the self-descriptiveness of each item—that is, they had distinct personality profiles (Figure 4a). On average, inter-participant similarity in self-descriptiveness ratings was significantly lower than that in semantic ratings (*t(495)* = –92.21, *p* < 0.001), indicating that how participants responded to each item during the self-reference task was more idiosyncratic than the semantic judgment task.

**Figure 4.**
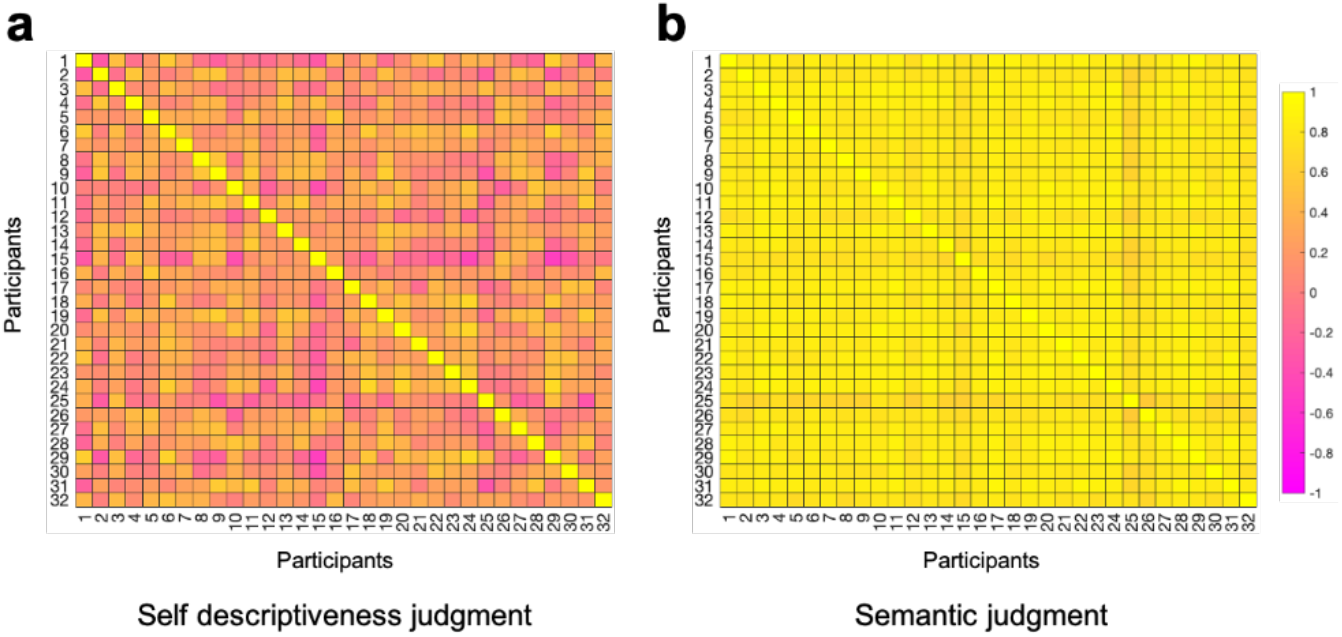
Similarity in self-descriptiveness judgment (**a**) and semantic judgment (**b**) across participants.

### fMRI results

#### Univariate analysis results

Replicating previous findings^5,6,37^, the contrast of self-reference versus semantic judgment significantly activated the medial prefrontal cortex (mPFC) (Figure 5a; × = –9, y = 47, z = –4; 344 voxels). No other brain regions showed significant activation for this contrast. It is worth noting that, in addition to the mPFC, prior studies have often reported significant activation in the posterior cingulate cortex (PCC) during self-referential processing. In the current study, two clusters were observed in the PCC (cluster 1: × = –9, y = –58, z = 20, 85 voxels; cluster 2: × = –6, y = –31, z = 44, 32 voxels) at an uncorrected height threshold of *p* < 0.005. However, neither cluster survived the cluster-level threshold of *p* < 0.05 (FWE-corrected). The reverse contrast (i.e., *semantic > self*) revealed significant activation in the bilateral ventrolateral prefrontal cortex (VLPFC) (left: × = –42, y = 41, z = –13, 207 voxels; right: × = 42, y = 41, z = –13, 148 voxels).

**Figure 5.**
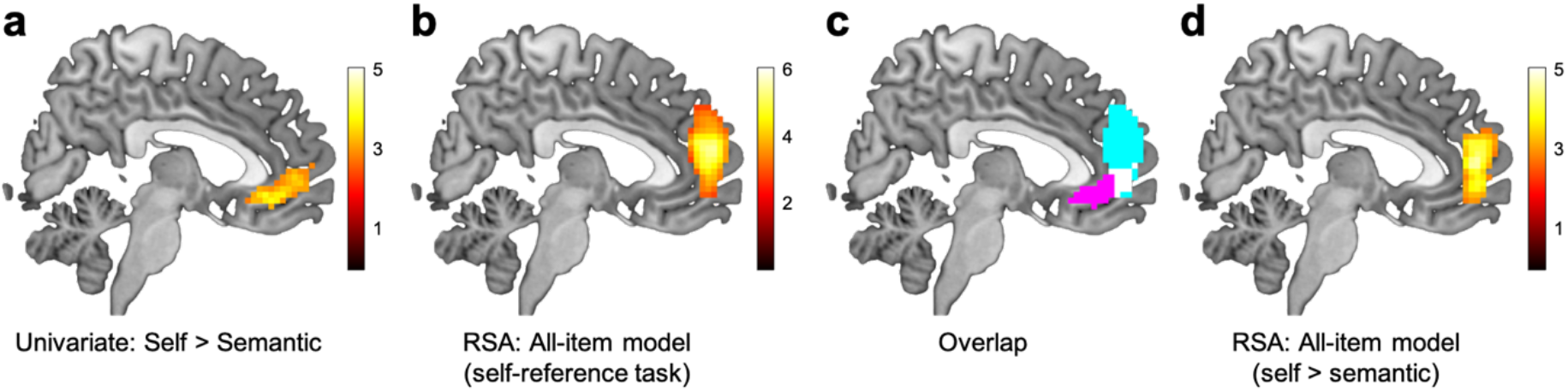
Results of the univariate analysis and RSA. (**a**) Univariate results showing regions significantly activated by the self > semantic contrast (x = 4). (**b**) RSA results for the all-item model, identifying regions where neural response similarity significantly depended on whether item pairs were from the same or different scales (x = 4). (**c**) Overlap between univariate and RSA results (x = 4). Magenta indicates activation from the univariate contrast (*self > semantic*), cyan indicates RSA results from the all-item model, and white indicates the overlapping region. (**d**) Results from a paired t-test comparing searchlight RSA maps from the self-reference and semantic judgment tasks (x = 4). mPFC neural response similarity reflected item scale membership significantly more during the self-reference task than during the semantic judgment task.

#### Searchlight RSA result

A searchlight RSA within the mPFC mask identified a significant mPFC cluster associated with the all-item model (Figure 5b; × = 3, y = 56, z = 17; 660 voxels), indicating that neural response similarity in this region varied depending on whether item pairs originated from the same or different psychological scales. Although this mPFC cluster overlapped with the mPFC area activated by the self > semantic contrast, the RSA-identified cluster tended to be located more dorsally than the univariate activation cluster (Figure 5c). Outside of the mPFC mask, no regions showed a significant association with the all-item model. Thus, the scale-dependent pattern similarity was found only in the mPFC.

In contrast, when the same searchlight RSA using the all-item model was applied to data from the semantic judgment task, no significant clusters were identified. A direct comparison between the two tasks (*self > semantic*) revealed a significant cluster in the mPFC (Figure 5d; × = 0, y = 50, z = 8; 372 voxels). No other significant regions were found. The reverse contrast (*semantic > self*) did not reveal any significant activation. These findings indicate that mPFC neural response patterns reflected whether items were from the same or different psychological scales, but only during the self-reference task. This suggests that the pattern similarity observed in the mPFC (Figure 5b & 5d) cannot be explained by low-level linguistic factors such as item meaning, grammatical structure, or lexical similarity, which were held constant across tasks.

#### ROI analysis results

##### RSA

We first confirmed that neural response similarity within the mPFC ROI (Figure 6a) during the self-reference task significantly reflected whether item pairs were from the same psychological scale, as predicted by the all-item model (*pperm* = 0.001; Figure 6b). In contrast, the effect was not significant for the semantic judgment task (*pperm* = 0.51), and the difference in model fit between the two tasks was itself significant (*pperm* = 0.008), replicating and extending the results observed in the searchlight analysis (Figure 5b & 5d).

**Figure 6.**
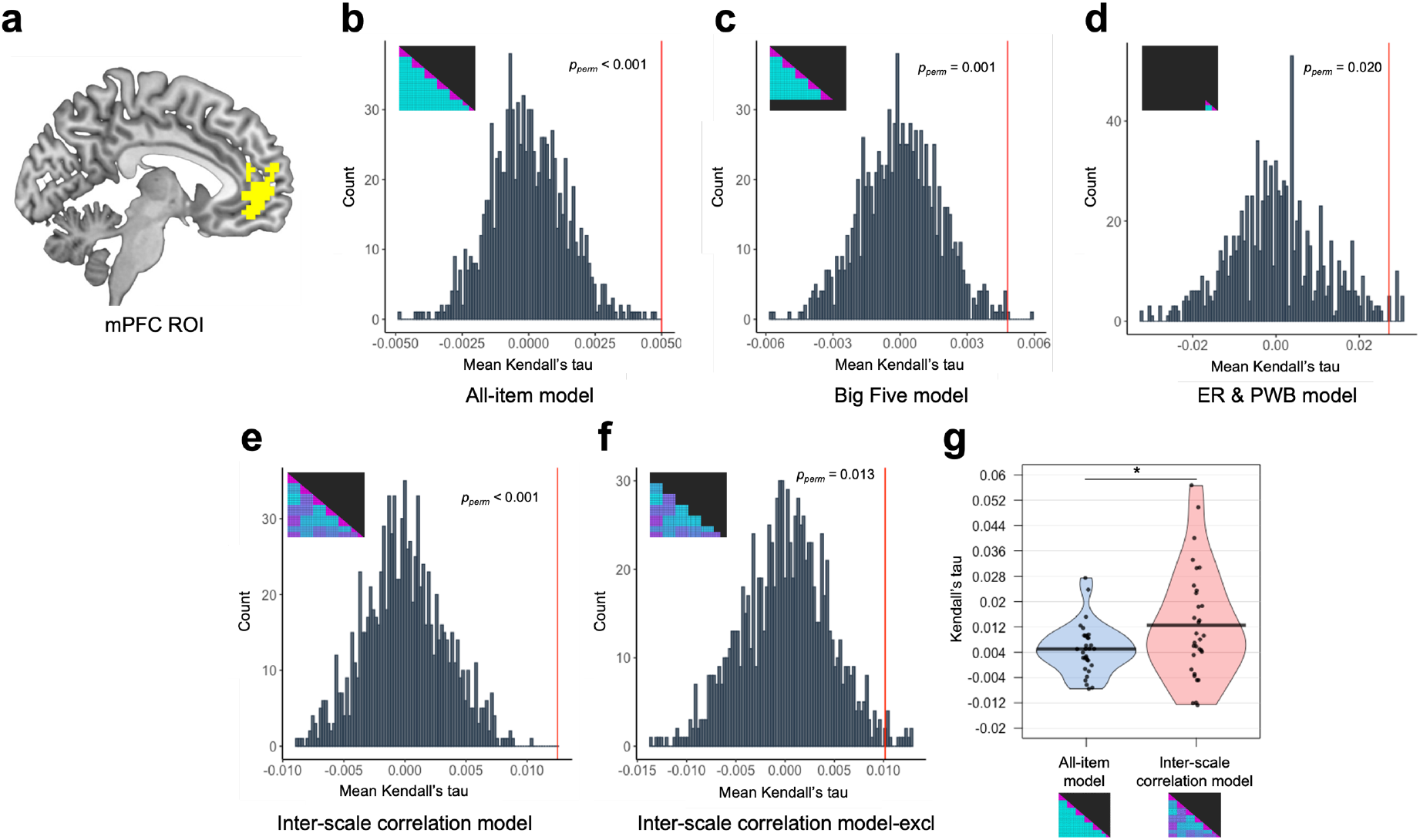
Results of the ROI analyses. (**a**) The medial prefrontal cortex (mPFC) ROI used in the analysis (x = –6), defined using a Neurosynth term-based meta-analysis for the term “self-referential.” (**b–f**) Results of permutation tests assessing the fit between neural RSMs and five model RSMs: (**b**) all-item model, (**c**) Big Five model, (**d**) PWB & ER model, (**e**) inter-scale correlation model, and (**f**) inter-scale correlation model–excl. (**g**) The inter-scale correlation model showed a significantly better fit to mPFC neural response patterns compared to the all-item model. * *p* < 0.05 (two-tailed Wilcoxon signed-rank test). Red vertical lines represent the mean Kendall’s tau correlation with the actual data.

We conducted the same analysis separately for the 60 Big Five items (Big Five model; Figure 2d) and the 12 items from the ER and PWB scales (ER & PWB model; Figure 2e) to assess whether the observed effect was primarily driven by the Big Five items. For both models, neural response similarity in the mPFC during the self-reference task was significantly associated with whether item pairs belonged to the same scale: the Big Five model (*pperm* = 0.001; Figure 6c), and the ER & PWB model (*pperm* = 0.020; Figure 6d). There was no significant difference in model fit between the two (*p* = 0.18, two-tailed Wilcoxon signed-rank test), suggesting that scale-dependent mPFC pattern similarity is not unique to the Big Five, but extends to other psychological constructs as well.

Thus far, our analyses focused on whether item pairs belonged to the same or different psychological scales. However, as shown in the behavioral results (Figure 3), the scales were themselves more or less correlated with one another. To test whether mPFC neural response similarity reflected the strength of inter-scale correlations, we constructed a model RSM based on the absolute values of correlation coefficients among the seven scales (inter-scale correlation model; Figure 2e) and computed its fit with the neural RSM. The results revealed a significant association between the two (*pperm* < 0.001; Figure 6e). To provide stronger evidence that mPFC neural similarity reflects graded psychological similarity, we conducted the same analysis using a modified version of the model that excluded all within-scale item pairs (inter-scale correlation model–excl; Figure 2f). Even with these within-scale cells removed, the association remained significant (*pperm* = 0.013; Figure 6f). Finally, we directly compared model fit between the all-item model and the inter-scale correlation model. The inter-scale correlation model provided a significantly better fit (*p* = 0.04, two-tailed Wilcoxon signed-rank test; Figure 6g). Taken together, these results indicate that mPFC neural response similarity reflected the degree of psychological similarity across scales, rather than categorical scale membership alone.

##### Multiple regression RSA: Controlling for semantic similarity across items

Some psychological scales have been criticized for including near-synonymous items, which may artificially inflate internal consistency without broadening the construct being measured^38^. It is therefore unsurprising that two near-synonymous items might elicit similar neural responses. Although the BFI-2^16^ was not explicitly developed to eliminate such redundancy, its item development emphasized clarity and trait breadth, and it appears to avoid obvious near-synonyms within each scale. Still, items measuring the same trait often exhibit semantic similarity, as they are designed to tap into related aspects of a common psychological dimension. Supporting this, a recent study using large language models found that Big Five items within the same trait domain are more semantically similar to one another than to items from other domains^39^. This raises the question of whether our observed neural similarity might reflect such semantic overlap rather than deeper psychological structure (— although a purely semantic account would not explain the mPFC specificity of our effects, nor their restriction to the self-reference task).

To directly address this issue, we conducted an additional analysis incorporating sentence-level semantic similarity, estimated using Sentence-BERT (Reimers & Gurevych, 2019). We constructed a semantic similarity RSM and tested whether the inter-scale correlation model explained mPFC neural similarity above and beyond semantic similarity (Figure 2g).

The results first showed that, consistent with the previous study^39^, within-scale semantic similarity was significantly higher than across-scale semantic similarity (*t*(2554) = 6.25, *p* < 0.001). Visual inspection of the similarity matrix suggested that semantic similarity for the six ER items was particularly high (see Figure 2g), but the difference between within- and across-scale similarity remained significant even when considering only Big Five items (*t*(1768) = 5.24, *p* < 0.001).

The results of the multiple regression RSA showed that the inter-scale correlation model remained a significant predictor of mPFC pattern similarity even after controlling for semantic similarity (mean *β* = 0.0024, *pperm* = 0.001). This suggests that multivoxel mPFC responses during self-referential evaluation reflect the psychological structure of the traits being assessed, rather than merely tracking surface-level linguistic similarity among items.

The semantic similarity model (based on Sentence-BERT) was also significantly associated with mPFC neural responses (mean *β* = 0.0041, *pperm* < 0.001), indicating that semantically similar items evoke similar activation patterns in the mPFC.

When we repeated the analysis using the semantic similarity model based on USE ^35^, the inter-scale correlation model again showed a significant association with mPFC neural responses (mean *β* = 0.0026, *pperm* = 0.001). In contrast, and unlike the result based on Sentence-BERT, the semantic similarity model derived from USE was not significantly associated with mPFC neural responses (mean *β* = 0.0010, *pperm* = 0.20).

## Discussion

The present study used RSA to investigate how the human brain—particularly the mPFC—represents psychological traits as assessed through common personality questionnaires. By combining multivariate fMRI analyses with theoretical and behaviorally derived models of psychological structures, we demonstrated that mPFC neural response patterns during self-referential judgments are sensitive to whether items are drawn from the same psychological scale (Figure 5b & 5d). No region other than the mPFC showed the same result. Importantly, this effect was not observed during semantic desirability judgments, indicating that our results cannot be explained by low-level semantic or linguistic features, as the same set of questionnaire items was used across the self-reference and semantic judgment tasks.

The ROI analyses further revealed that this effect was not limited to a particular set of traits. Both Big Five items and items from emotion regulation and psychological well-being scales showed scale-sensitive pattern similarity in the mPFC, with no significant difference between them (Figure 6c & 6d). Moreover, we found that mPFC similarity patterns were not merely categorical (same vs. different scale) but reflected graded psychological similarity, as indexed by inter-scale correlations (Figure 6e). To further rule out the possibility that these results were driven by sentence-level semantic similarity, we conducted a multiple regression RSA including a semantic similarity model (estimated using Sentence-BERT^40^ and USE^35^) as a covariate. Critically, the inter-scale correlation model remained a significant predictor of mPFC neural similarity even after controlling for semantic similarity, suggesting that the mPFC encodes psychologically meaningful trait structure beyond surface-level linguistic similarity.

The pattern of results observed in the mPFC likely reflects the underlying psychological similarity in how individuals respond to each scale item. Specifically, each item from a well-constructed psychological scale prompts respondents to access similar aspects of their self-concept—potentially represented in the mPFC in terms of self-importance (see ^25^)—and/or to retrieve relevant information in a similar manner to form a judgment and select a response (e.g., drawing on similar personal memories and contextual self-knowledge)^14^. This process, in turn, results in similar activation patterns in the mPFC across items belonging to the same scale. Our previous study^14^ demonstrated that mPFC neural responses during a self-reference task with trait adjectives reflect multiple cognitive processes, some of which are shared with other tasks involving judgments based on internally constructed representations (e.g., other-reference, introspection, and autobiographical memory tasks). The present study extends this work by showing that even within the same self-reference task, mPFC responses distinguish among items from different psychological scales. Crucially, our multiple regression RSA demonstrated that this scale-sensitive pattern similarity in the mPFC could not be explained by surface-level linguistic similarity alone. Instead, it reflects the underlying psychological structure of the traits being assessed.

Thus, the present findings highlight the representational richness of the mPFC in encoding fine-grained aspects of self-relevance. Previous multivariate neuroimaging studies have shown that activation patterns in the mPFC differentiate between self- and other-referential processing^41-44^. Feng et al.^44^ further demonstrated that mPFC activation patterns during the self-reference task vary depending on the type of self-knowledge being accessed— for example, traits, physical attributes, or social roles. The present study builds on this work by showing that mPFC patterns not only distinguish between items from distinct psychological constructs within standard personality questionnaires, but also reflect graded psychological similarity across scales—suggesting a representational structure that goes beyond categorical differences.

The present findings also contribute to a theoretical understanding of self-report not merely as a measurement tool but as a cognitive process in its own right. Our results suggest that responding to a questionnaire item involves more than retrieving a static belief or trait; instead, it engages an active construction of self-relevant evidence, supported by integrative processes in the mPFC. This view aligns with models of judgment that emphasize the dynamic nature of introspection and decision-making^2^, and extends them by providing neural evidence that different items—depending on their psychological content—invoke distinct yet systematically related patterns of internal processing. From this perspective, self-report can be seen as the behavioral outcome of a representational evaluation process, in which the brain draws upon personal knowledge, situational context, and normative expectations to produce a coherent response. Understanding self-report in these terms has implications for how we interpret questionnaire data—not as direct reflections of latent traits, but as outputs shaped by structured, construct-specific mental operations.

Moreover, our study contributes to the growing interest in functional dissociations between the mPFC and PCC within the default mode network. The absence of significant effects in the PCC in the present study, despite its frequent involvement in self-referential processing^37^, highlights the specificity of the mPFC’s role in structured psychological evaluation. The PCC has been implicated more strongly in autobiographical memory retrieval and self-projection into imagined events^10^ (see also ^13^), whereas the present task emphasized abstract, conceptual judgments about stable traits. Our findings suggest that the mPFC, rather than the PCC, serves as the principal site for integrating internally constructed evidence when individuals evaluate themselves in a questionnaire context.

Finally, our results suggest an intriguing avenue for future research at the intersection of cognitive neuroscience and personality psychology: the possibility of using neuroimaging methods to assess the validity of psychological scales. While there are well-established procedures for scale development and validation^45^, these procedures rely almost entirely on subjective sources of evidence—such as expert judgment and participants’ self-reports. The present findings raise the possibility that fMRI could provide objective and complementary information about whether a scale measures a coherent psychological construct at the neural level. That said, given the relatively small effect sizes observed in the present study (i.e., the magnitude of neural–model RSM correlations; see Figure 6), such an approach should currently be viewed as a supplement—not a replacement—for established psychometric practices. Neuroimaging-based validation remains limited by practical constraints, including issues of scalability, cost, and interpretability. Nonetheless, our findings suggest that when questionnaire items are well-constructed to target a common construct, their neural representations—as reflected in mPFC activity—may mirror the same psychological structure observed in behavioral responses. Incorporating neural data into scale validation efforts could eventually help address foundational challenges in personality research^46^ and contribute to efforts to defragment psychology^47^.

In sum, our findings provide new evidence that neural responses in the mPFC reflect the psychological constructs targeted by self-report questionnaires. By applying RSA, we demonstrated that the mPFC encodes both categorical and graded similarities among trait constructs—but only when individuals evaluate themselves. These results shed light on how the brain organizes self-knowledge and highlight a promising link between neuroscience and personality assessment. Given the central role of self-report across many areas of research, understanding its neural basis may help strengthen the bridge between subjective experience and objective measurement.

## Acknowledgements

We thank Haruto Yaegashi and Ryuji Saito for their assistance with fMRI data collection. This research was supported by a Japan Society for Promotion of Science (JSPS) KAKENHI Grant Number JP19K24680 (to K.I.).

